# Identifying Neural Signatures of Dopamine Signaling with Machine Learning

**DOI:** 10.1101/2023.01.17.524454

**Authors:** Siamak K. Sorooshyari, Nicholas Ouassil, Sarah J. Yang, Markita P. Landry

## Abstract

The emergence of new tools to image neurotransmitters, neuromodulators, and neuropeptides has transformed our understanding of the role of neurochemistry in brain development and cognition, yet analysis of this new dimension of neurobiological information remains challenging. Here, we image dopamine modulation in striatal brain tissue slices with near infrared catecholamine nanosensors (nIRCat) and implement machine learning to determine which features of dopamine modulation are unique to changes in stimulation strength, and to different neuroanatomical regions. We trained a support vector machine and a random forest classifier to determine whether recordings were made from the dorsolateral striatum (DLS) versus the dorsomedial striatum (DMS) and find that machine learning is able to accurately distinguish dopamine release that occurs in DLS from that occurring in DMS in a manner unachievable with canonical statistical analysis. Furthermore, our analysis determines that dopamine modulatory signals including the number of unique dopamine release sites and peak dopamine released per stimulation event are most predictive of neuroanatomy yet note that integrated neuromodulator amount is the conventional metric currently used to monitor neuromodulation in animal studies. Lastly, our study finds that machine learning discrimination of different stimulation strengths or neuroanatomical regions is only possible in adult animals, suggesting a high degree of variability in dopamine modulatory kinetics during animal development. Our study highlights that machine learning could become a broadly-utilized tool to differentiate between neuroanatomical regions, or between neurotypical and disease states, with features not detectable by conventional statistical analysis.

## Introduction

Recent advances in the ability to image neuromodulators from single neurons (1), in acute brain slices (2) and in vivo (3), (4) have enabled insights into the role of neurochemical communication in the neurotypical and diseased brain. These newly accessible neurochemical signals are poised to revolutionize neuroimaging by providing an additional dimension of information regarding the role of neuromodulation in regulating brain circuits and the central role of neuromodulators in psychiatric and neurodegenerative disease. As these neuroimaging tools are implemented more broadly, it becomes imperative to interpret the biological underpinnings of these neurochemical signals and understand the features of importance of this previously ‘invisible’ dimension of neurobiology. Herein, we implement several machine learning approaches to identify and evaluate what features extracted from dopamine imaging studies are important for distinguishing neurobiological features of importance, and which machine learning approaches enable these analyses.

Male B6CBA-Tg(HDexon1)62Gpb/3J mice (R6/2 mice) were purchased from Jackson Labs and bred at 6 weeks with 10 week old female C57BL/6 mice. Near-infrared catecholamine nanosensors (nIRCat) have enabled imaging of neuromodulator dopamine in brain tissue (2) and to study the role of its modulation in neurodegeneration (17). nIRCat is a versatile synthetic optical tool for monitoring dopamine release and reuptake in acute slices and is compatible with a broad range of pharmacological agents used to target dopamine receptor activity. nIRCat provides high spatial (micron) and temporal (second) resolution videos of dopamine modulation in the brain extracellular space, with many features contributing to the signatures of dopamine release and reuptake through its volume transmission in the extracellular space, enabling time-resolved imaging of dopamine modulation at the level of individual synapses. In this work, we use nIRCat to image electrically-stimulated dopamine release within the dorsal striatum of acute brain slices generated from wildtype 4, 8.5, and 12 week old mice. Moreover, we assess whether machine learning can be implemented to identify whether dopamine modulatory signals can distinguish striatal subregions, and which features of neurotransmitter modulation are most predictive to identify different brain regions. Specifically, we apply two conventional yet distinct machine learning techniques to analyze stimulated dopamine release imaged with nIRCat in the dorsolateral striatum (DLS) and the dorsomedial striatum (DMS). Stimulated dopamine release is achieved with a single pulse at either 0.1 mA or 0.3 mA stimulation strength. The two machine learning approaches we implemented are a support vector machine (SVM) and, separately, a random forest (RF) approach. We selected a SVM because of its capability to distinguish observations into separate classes based on features that may share complex, nonlinear relationships with the different classes to which the observations belong. The SVM notion is based on finding a boundary in a space that separates the training data into distinct classes and then applying that same boundary or rule to the test data. We selected the RF approach because it is a relatively simple method that will allow interpretation of which variables from our dopamine imaging datasets enable the most accurate predictions. RF is an ensemble method where multiple decision trees are formed on the same dataset. The individual decisions made from each decision tree are then combined to arrive at a consensus on the outcome that is output as the classification decision. The classification decision is output after the decision criteria for each split at a node of a tree are determined, based on the training data in supervised fashion. The intricacy of RF that distinguishes it from its decision tree relatives is that RF decorrelates the decision trees by considering only a subset of features at each split in a constituent tree. This approach prevents the possibility of one or a small portion of the features dominating the decisions.

In our experimental workflow, we stimulated nIRCat-labeled acute striatal slices with a single 0.1 mA or 0.3 mA pulse and imaged dopamine release in the DLS or DMS striatal regions of mouse brain tissue. Our hypothesis is that any differences in dopamine release in DLS versus DMS could be elucidated from our nIRCat data features, and that machine learning approaches can unearth whether dopamine imaging is alone sufficient to distinguish the DLS from DMS. Recent work has shown there exist differences in signaling, specifically differences in peak dopamine concentration between dorsal and ventral subregions of the striatum in mice. In (15) Calipari et al. found that DLS produces a roughly 4x higher concentration of stimulated dopamine release than the Nucleus Accumbens core when measured in acute brain slices with fast-scan cyclic voltammetry (15). However, somewhat contradictory results have been reported using R6/2 mice, in which no significant differences in maximum dopamine release were observed across striatal subregions (16). Known differences in dopamine transporter levels across the dorsal striatum indicate that peak concentration may not fully capture signaling differences in the basal ganglia, and indeed that regional differences in autoreceptor expression, differences in axonal architectures, or differences in projection density may contribute to signaling kinetics (20), (21), (22). We therefore sought to study what dopamine signaling features contribute to regional differences in dopamine modulatory kinetics and assess whether machine learning approaches may provide a user-removed approach to identifying subtle changes between brain regions in a manner that considers more than absolute dopamine release concentrations. We further hypothesize that machine learning can identify which features of dopamine modulation enable this differentiation. We anticipate that identification of features of stimulated dopamine release to be important for differentiating neuroanatomy and confirming the role of machine learning for this goal, will provide important insights for interpretation of neuromodulator imaging experiments broadly.

## Results

### Collected data

Male B6CBA-Tg(HDexon1)62Gpb/3J mice (R6/2 mice) were purchased from Jackson Labs, which we bred at 6 weeks with 10 week old female C57BL/6 mice. Mice were kept in temperature-controlled environments with three to five mice per cage on a 12-hour light/dark cycle, with all animal procedures approved by the University of California Berkeley Animal Care and Use Committee (ACUC). We synthesized our dopamine nIRCat nanosensor and used nIRCat to label acute live brain slices as described previously in (Yang et al., 2021). Acute brain slices were then labeled with nIRCat through passive incubation in 5 ml of ACSF containing nIRCat nanosensor at a concentration of 2 mg/L for 15 minutes, rinsed, and placed in a recording chamber to equilibrate during which a tungsten bipolar stimulation electrode was positioned in the dorsal-lateral striatum. We applied a single electrical stimulation pulse of 0.1 mA or 0.3 mA after collecting 200 frames of baseline nIRCat fluorescence from two brain regions: DLS and DMS. To reduce bias, all stimulation videos were collected in triplicate and we alternated stimulation strengths. Next, nIRCat-labeled brain tissue slices were processed to quantify their response to stimulated dopamine release. Raw Image stack files were processed using a custom-built, publicly available MATLAB program (https://github.com/jtdbod/Nanosensor-Imaging-App) with the image processing protocol described in depth elsewhere (18). These processed data form the inputs of our machine learning algorithms, summarized in Table 2, and are calculated as follows: Regions of dopamine release in acute slice tissue are identified by large changes in nIRCat ΔF/F response. Dopamine hotspots were programmatically identified by creating a grid of 2 μm squares across the field of view to reduce bias and expedite image stack processing time. The relationship (F - F0)/F0 was used to calculate the change in fluorescence, ΔF/F, of each grid square. Herein, F0 represents the average fluorescence of the grid square over the first 30 frames of the image stack and F is the fluorescence intensity of the grid square as it changes over the 600 collected frames. The program identifies dopamine hotspots as regions of interest with statistically significant dopamine release activity if these grid squares exhibit nIRCat fluorescence behavior that is at least 3 standard deviations above the baseline fluorescence activity, F0, at the time of slice stimulation (200 frames). In this manner, we identified dopamine release hotspots for each stimulation image stack for nIRCat-labeled brain slices. The peak fluorescence, ΔF/F, of each dopamine hotspot in the image stack was averaged to generate the average image stack peak ΔF/F. The number of active dopamine release sites per stimulation event per brain slice were then identified and averaged to output the average slice hotspot number. Mean values for dopamine release and reuptake curves were calculated from averaging traces from each slice, including 3 stimulations per slice and 1 slice per mouse. These data outputs were subsequently used herein to assess the feasibility of machine learning to distinguish differences in features, such as peak ΔF/F or dopamine hotspot number, across stimulation strengths and across brain regions. Table 1 lists the experimental conditions and number of independent stimulations used for each experimental condition, and Table 2 lists the collected and analyzed data from nIRCat-labeled brain slices subjected to single-pulse stimulation for dopamine release.

**Table 1.**
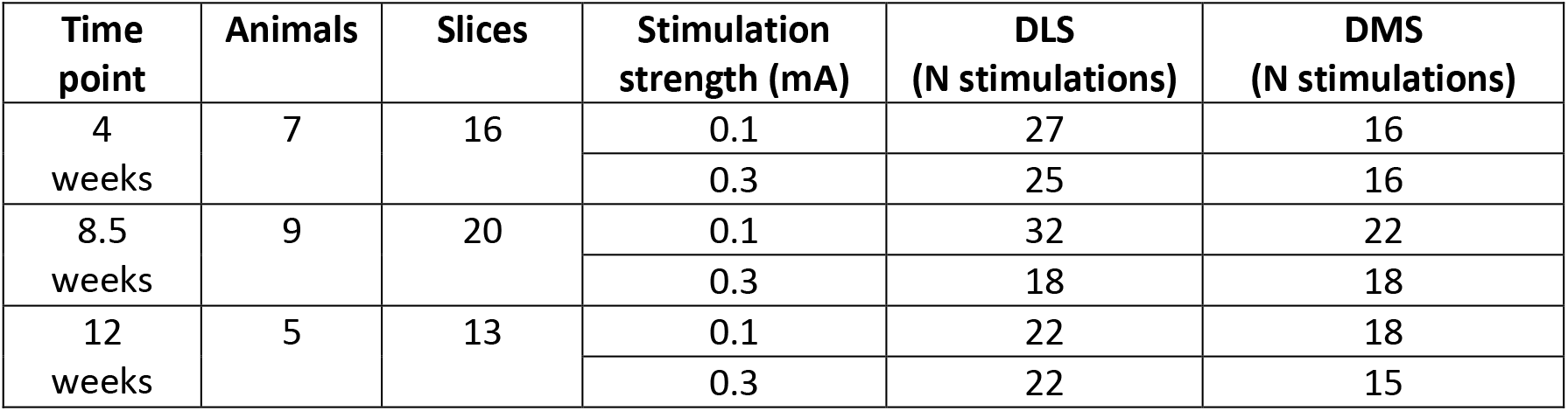
Distribution of brain regions and stimulation strengths that were recorded in the brain slices of WT mice.

**Table 2.** The features computed from traces to be used in the machine learning analysis. Features were calculated for each recording and used as the inputs to the algorithms.

| Feature | Description |
| --- | --- |
| E[dF] | mean of dopamine dF trace |
| Var[dF] | variance of dopamine dF trace |
| med[dF] | median of dopamine dF trace |
| min[dF] | Minimum dopamine dF trace |
| max[dF] | Maximum dopamine dF value in recording |
| ROI count | number of dopamine release sites (ROIs) in the brain slice field of view |
| AUC of dF | area under the dopamine dF curve |
| decay of dF | Dopamine reuptake kinetics calculated by an exponential decay fit to the dopamine dF trace |
| ND-low | number of times dF was lower than the $-2\sigma$ level |
| ND-high | number of times dF exceeded the $2\sigma$ level |

The first 8 features (red) are deemed as statistical features whereas the last 2 features (blue) are referred to as paroxysmal features. In several of the ML analyses, all 10 features were simultaneously used.

### Computed features

Acute coronal brain slices were generated to contain the dorsal striatum and labeled with nIRCat nanosensor (Figure 1A) as described previously (2) to image stimulated dopamine release as described above and schematically depicted in Figure 1(B) and (C). Time series traces were used to generate the data features listed in Table 2, and prior to this study it was unknown which of these features would be most informative for distinguishing DLS from DMS and indeed for analysis of neurochemical modulation data as a whole (Figure 1D). Table 2 summarizes the features that were analyzed from each brain slice labeled with nIRCat and following electrical stimulation. We also provide a representative intensity trace annotated to highlight features used in the present study (Figure 1D) and are as follows: The number of active putative dopamine release sites, termed regions of interest (ROIs), are programmatically identified as described previously (2) and represent dopamine release sites. Each individual ROI in turn includes a series of features associated with dopamine release and reuptake from a single dopamine release site, which includes maximum change in fluorescence (max dF), area under the dopamine response curve (AUC), and dopamine signal decay (τ). Max dF is proportional to the maximum concentration of dopamine measured by nIRCat at that release site. AUC represents the integrated area under the ROI fluorescence curve following stimulated release and is a relative measurement of the total amount of dopamine released by a release site. Signal decay represents the reuptake kinetics of dopamine, a signal that is dependent on the expression and activity of dopamine transporters in brain tissue within the ROI. Each of these features are computed over the duration of each recording to attain the mean, variance, median, minimum, and maximum values, where the last two features in Table 2 reflect the number of “deviations” for a recording, termed paroxysmal features. Specifically, each stimulated brain slice was Z-scored and a deviation-high was declared when the recording exceeded a value of 2 standard deviations from the mean, similarly a deviation-low was declared when the recording fell below a value of −2 standard deviations from the mean. The number of low and high deviations in a recording are referred to as ND-low (ND: number of deviations) and ND-high, respectively. In training machine learning algorithms and making ensuing predictions, we consider a combination of all 10 features from Table 2 as the combined feature set.

**Figure 1.**
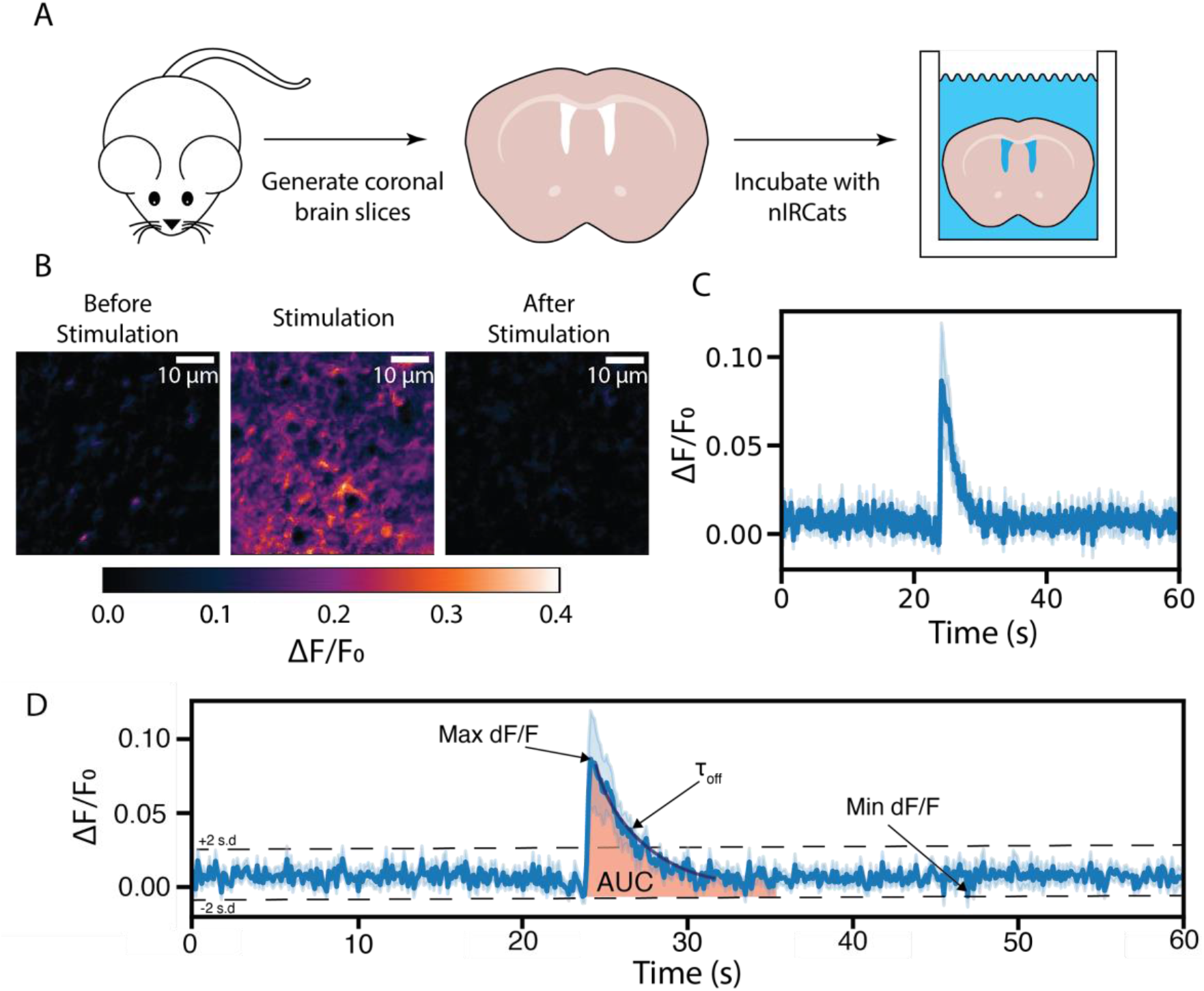
The feature extraction procedure of using nIRCat to image electrically-stimulated dopamine release in mouse brain slides. (A) Cartoon depiction of generation of nIRCat-labeled brain slices. (B) Representative images of stimulated dopamine release in acute striatal slices. (C) Processed dopamine trace depicting release of dopamine after electrical stimulation from a single dopamine release site. (D) A single dopamine trace annotated with mined features.

### Machine learning algorithm development

The SVM and RF algorithms were trained on the features in Table 2, first to assess whether these algorithms could differentiate between dopamine released from nIRCat-labeled slices stimulated at either 0.1 mA or 0.3 mA stimulation strengths. Prior literature using nIRCat labeled brain slices shows that different stimulation strengths clearly generate proportionately different levels of dopamine release, with higher stimulation amplitudes generating a higher max dF/F and AUC (2). Therefore, we trained our algorithms on these data for which the dopamine release differences are known. The SVM algorithm used a linear kernel with a binary classifier. With the kernel denoting a function that reflects the similarity among observations - the use of a linear kernel leads to the decision boundary between the classifier’s decisions being a linear function of the considered features. The choice of a binary classifier was made because in each prediction we are considering two alternatives, i.e. distinguishing between DLS/DMS regions or distinguishing between 0.1/0.3 mA stimulation strength. For the predictions that are made in our analysis, the linear kernel was selected for its simplicity and the absence of detailed *a priori* knowledge regarding which of the dopamine features would be most important in distinguishing between stimulation strengths and eventually between brain regions from which dopamine originates. We did not consider the RF algorithm with paroxysmal features as there were too few features (i.e p = 2) to consider this as worthwhile analysis. We next evaluated the predictive capability with the SVM and RF algorithms via a leave-one-out analysis with Monte-Carlo sampling of all animals and brain slices. The Monte-Carlo analysis consists of the data being repeatedly divided into a training and test set with the test data consisting of a single observation while the remaining data was evenly partitioned into two groups and used to train the machine. The computational pipeline is depicted in Figure 2 and was used to make predictions from the considered data.

**Figure 2.**
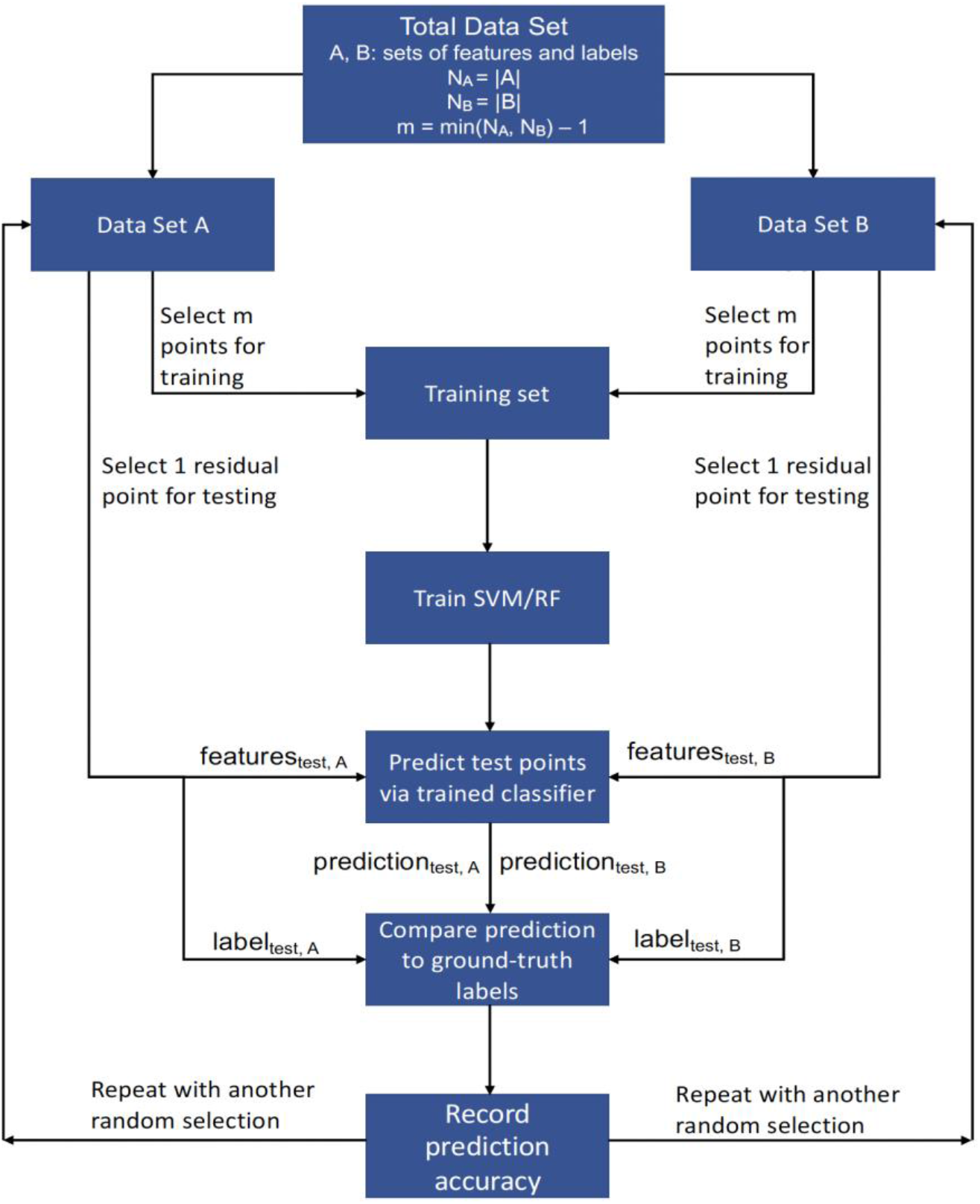
Machine learning workflow to predict the electrical stimulation strength applied to brain slices or the brain regions of dopamine release. The sets A and B may correspond to the groups of brain slices stimulated at 0.1 or 0.3 mA – alternatively, the sets may correspond to the groups of brain slices stimulated in the DMS or DLS. The notation |.| refers to the number of elements in the set (i.e. cardinality). The above sequence is repeated 1000 times to arrive at a classification accuracy rate (AR). Each iteration encompasses a SVM or RF being trained on features from training data prior to the machine being presented with one left-out data point from each of the two groups. The AR is attained by computing the fraction of times the labels of the left-out samples from set A and B were correctly predicted as being equal to their ground-truth values.

### Machine learning can distinguish dopamine release at different stimulation strengths

Stimulation of nIRCat-labeled brain slices shows higher dopamine release as a function of stimulation strength, as expected and as previously demonstrated (2). Therefore, to validate the ability of our machine learning algorithm to distinguish between dopamine modulatory behavior with a known dependence on experimental condition, we first assessed whether our algorithms could distinguish between nIRCat-labeled brain slices stimulated with either 0.1 mA or 0.3 mA stimulation strengths. To avoid the confounding effect of brain region on this analysis, we compared 0.1 mA versus 0.3 mA stimulation in DLS separately from DMS.

The SVM and RF algorithms were implemented with the different groups of features listed in Table 2, running SVM with all 10 features is denoted via SVM (10), while using only the 8 statistical and 2 paroxysmal features is denoted by SVM (8) and SVM (2), respectively. The same notation is used with the RF algorithm - i.e. RF (10) represents implementing RF with all 10 features. The pipeline in Figure 2 was followed in each case to attain an accuracy rate for each of the stimulation strengths. We use notation such as SVM (10) and RF (8) to denote the use of SVM with the 10 combined features - i.e. 8 statistical and 2 paroxysmal - and the use of RF with the 8 statistical features, respectively. Our results indicate that an accurate discernment of the stimulation strength in the DLS and DMS is not possible from brain slices generated from 4 and 8.5 week old mice with neither SVM or RF algorithms, although SVM performed slightly better in DLS than in DMS at 4 weeks. At 4 weeks, the aggregate accuracy (“aggregate” denotes averaging the 0.1 mA and 0.3 mA results to provide a holistic account) in the DMS does not exceed chance, while the best discernibility occurs from data taken in the DLS with the SVM algorithm and is 0.557 (Figure 3A). At 8.5 weeks, nIRCat features provide a slightly better indication for whether a 0.1mA or 0.3mA stimulation amplitude was used in the DMS region. Here, the aggregate accuracy in both DMS and DLS consistently but barely exceeds chance, with the best accuracy rate being 0.649 for the RF (8) algorithm (Figure 3B). However, as mice age to 12 weeks, the distinguishability generally increases when using the SVM or RF algorithms in both the DLS and the DMS brain regions. At 12 weeks, the aggregate accuracy in both DLS and DMS consistently exceeds chance for both algorithms with the best discernability occurring from data taken in the DLS with the RF algorithm at an accuracy rate of 0.832 (Figure 3C). It is interesting that the prediction accuracy of our machine learning algorithms consistently increase as a function of animal age, going from chance at 4 weeks to maximum accuracy rates of 0.649 (RF (8)) at 8.5 weeks and 0.78 (SVM (10)) at 12 weeks. Biologically, a 4 week old mouse represents a prepubescent young animal, with the 8.5 week timepoint representing mice that have shifted into adulthood and sexual maturity, and 12 week old mice well into adulthood. Our results confirm that dopamine dynamics are still developing across our timepoints, with much more variability in the kinetics and features of dopamine release in young animals that confounds our machine learning algorithms and prevents distinguishing stimulation strengths in 4 (and to a lesser extent 8.5) week old animals. Interestingly, by the time animals age into adulthood by 12 weeks, it is possible that dopamine release and reuptake features have stabilized and enable us to clearly distinguish between stimulation strengths, and indeed represents the adult age group of prior work showing increasing dopamine release as a function of stimulation strength (2). These results are important, because they suggest that dopamine measurements taken in prepubescent animals undergoing development may introduce a high degree of biological variability and may prevent experimentation using dopamine signaling outputs as the sole form of measurement, particularly when the predicted biological effect size is small or moderate. Conversely, dopamine imaging measurements taken from adult animals exhibit less biological variability, enabling our machine learning algorithms to clearly distinguish between stimulation amplitudes used, as expected.

**Figure 3.**
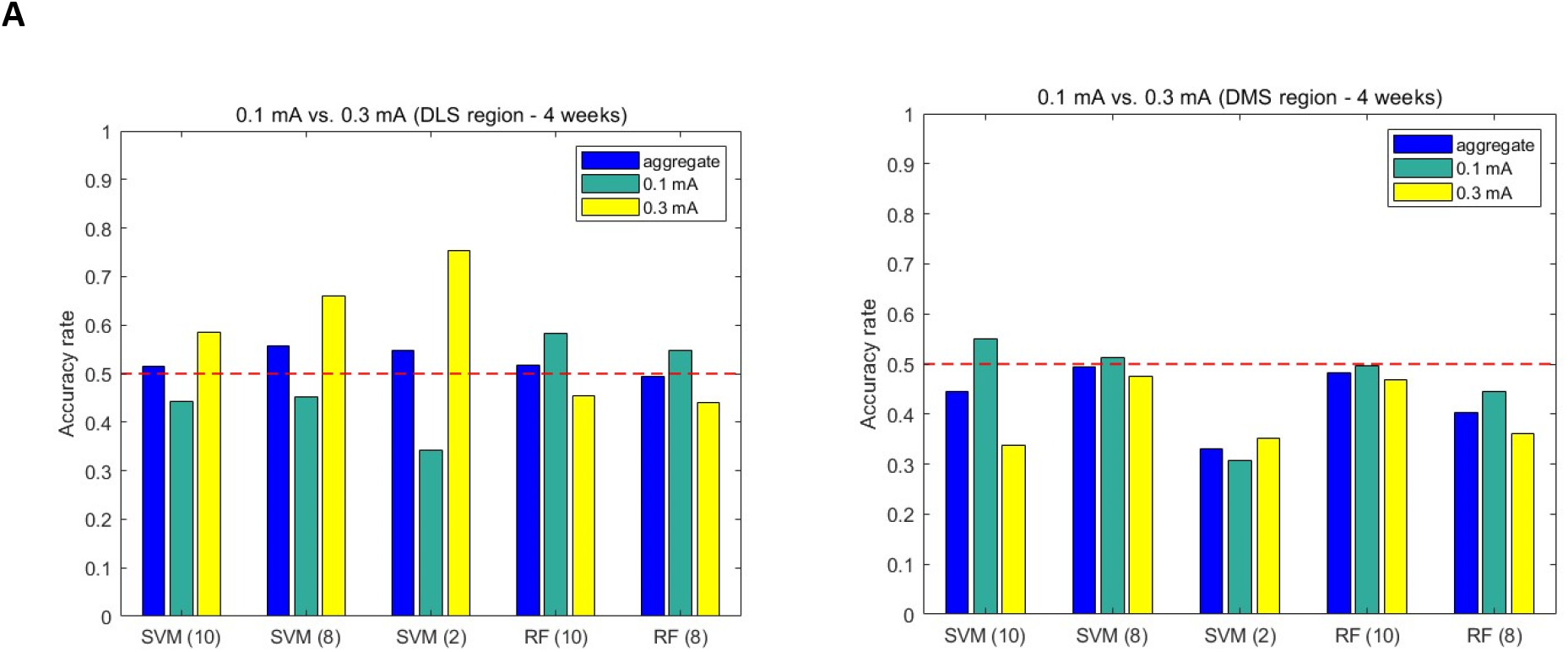
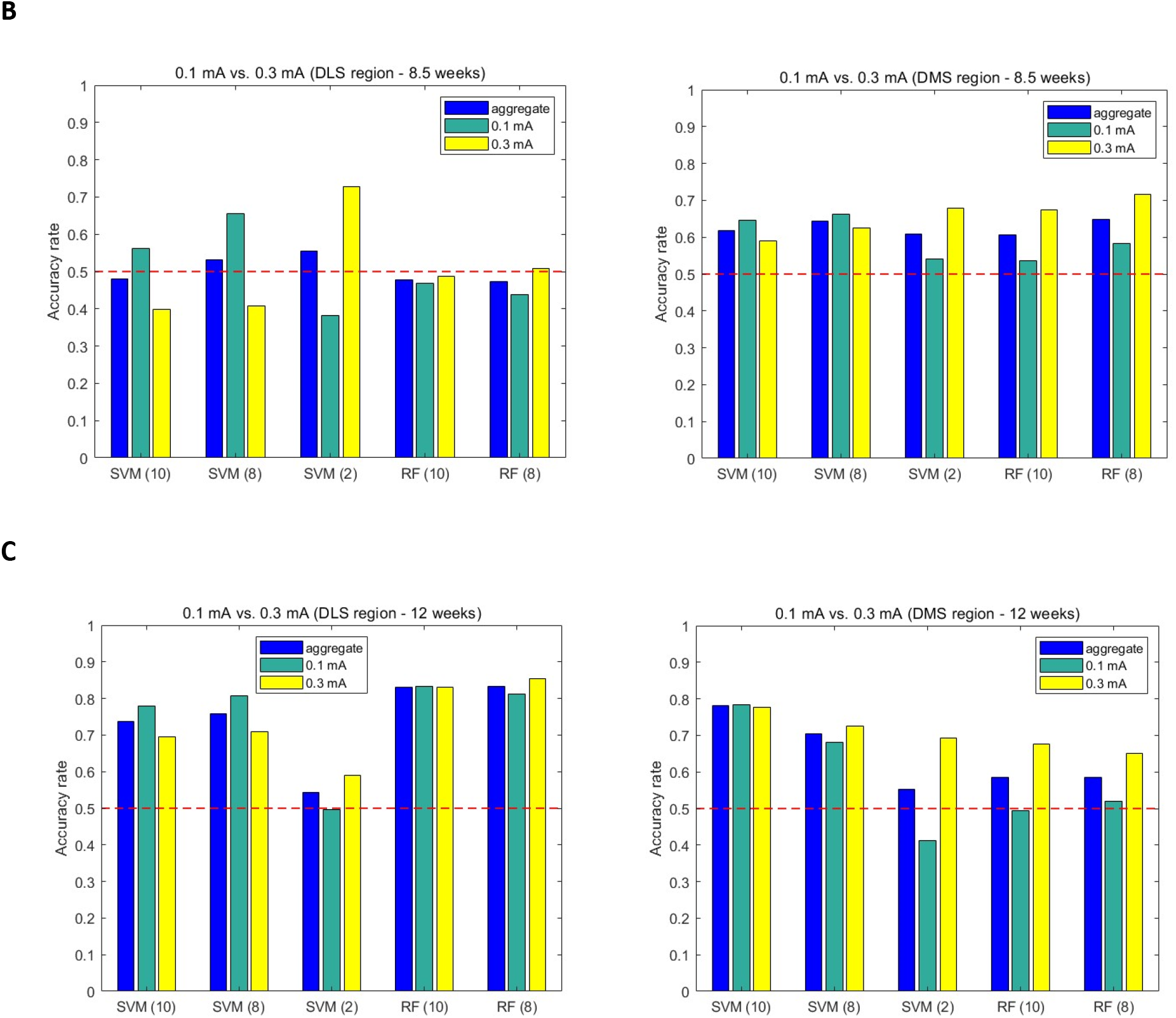
The predictive accuracy in distinguishing between 0.1mA and 0.3mA stimulation strengths of the DLS and DMS. Different time points were considered in the ages of the mice - subfigures (A), (B), and (C) correspond to results for the data collected from animals of 4, 8.5, and 12 weeks of age, respectively. Maximal predictive capabilities range from chance, 0.649, to 0.78 as a function of mouse age, respectively.

### Machine learning algorithm implemented to explore dopamine modulatory differences between DLS and DMS brain regions

Having confirmed that our SVM and RF algorithms are able to distinguish between brain tissue slices stimulated at 0.1 mA or 0.3 mA stimulation strengths in adult mice, as expected, we next sought to learn whether machine learning could be used to distinguish DLS or DMS striatal brain regions based only on the dopamine release and reuptake features from each region. To this end, we implemented the Monte Carlo technique with 1000 iterations for a single brain recording randomly selected from the DLS or DMS at each iteration to train our SVM and RF algorithms. To use a balanced training set for the predictive analysis, an equal number of DLS and DMS brain slices were used to train the machine prior to testing on the one held-out DLS and DMS brain slices, which represents a recording from the DLS region that was not used as part of the training data in distinguishing DLS from DMS. The accuracy rate in predicting whether a recording was made from the true striatal brain region was evaluated by counting the number of times that a correct classification was made for the held-out DLS and DMS samples across all iterations. The result is normalized by the number of iterations, 1000, used as part of the machine training pipeline, and an aggregate accuracy rate was computed by averaging the accuracy rate of held-out data from the DLS with that attained with held-out data from the DMS. We consider the predictive capability of our algorithm as being acceptable if the two constituent accuracy rates exceed the chance value of 0.5. Chance is a sufficient cutoff value since expert experimenters are unable to identify any of the tested classification problems better than the chance rate. The steps for the analysis and the prediction on test data are depicted in Figure 2, and the machine learning pipeline is the same as we used to assess whether the stimulation strength - i.e. 0.1 mA or 0.3 mA - can be determined from the dopamine modulatory signals.

It is often beneficial to be cognizant of the features that are the most important in making a classification, whether it be to distinguish stimulation strength or brain region. The RF machine learning algorithm is a tree-based method that readily provides an account of the features that are the most important in arriving at its decisions. Biologically, it is possible that certain features may be more important in distinguishing stimulation strengths or brain regions - e.g., to distinguish between stimulation strengths, a higher stimulation amplitude may evidently increase the max dF/F (more dopamine) without affecting the decay of dF (the rate at which dopamine is cleared from tissue). Conversely, differences in dopamine transporter expression in DLS versus DMS may affect dopamine clearance rates from tissue without affecting dopamine release features. For the development of our RF algorithm, we therefore utilized a node purity metric to evaluate which of the stimulated dopamine features are the most important to the RF techniques/decisions. For RFs, a node purity metric is equated to the total variance computed across the classes, with lower variance levels associated to a feature implying that in the trees’ arriving at a decision, a decision based on that feature contained a high percentage of data from one class (i.e. 0.1 mA vs 0.3 mA or DLS vs. DMS). Thus, the node purity metric provides a quantitative measure of the features’ importance in the RF classification decision. Unlike the RF algorithm that accounts for the relative contribution of each feature in arriving at its decision, the SVM algorithm does not. For SVM, the solution is a vector that determines a hyperplane setting the boundary between the decision regions. The vector for SVM is the solution of a quadratic optimization problem, and the variable section is typically applied by penalizing a norm on the optimization vector to suppress unimportant features from appearing in the solution. An alternative brute-force means of performing feature selection for SVM involves iteratively removing features or groups of features and evaluating the accuracy of the procedure on portions of the data set.

### Differentiating dopamine release from DLS versus DMS with nIRCat recordings

We next used the machine learning pipeline developed to differentiate dopamine dynamics to study nIRCat recordings made in the DLS or DMS of acute brain slices collected from 4, 8.5, or 12-week old mice as depicted in Figure 2. For this approach, we used data only from 0.3 mA stimulations, as they provided clearer nIRCat signals than the lower 0.1 mA stimulation amplitudes. SVM and RF approaches were used to process stimulated dopamine imaging videos on datasets segregated by animal age. The results shown in Figure 4 indicate that at 4 weeks of age, SVM provided better ability to distinguish whether dopamine was imaged from the DLS or DMS of a brain slice: an SVM operating on the 8 statistical features as listed in Table 2 provided the best performance via an accuracy rate of 0.615. RF also provided above-chance predictive capability with an accuracy rate of 0.573 and 0.569 for the combined and statistical features, respectively, but inferior to that attained with the SVM for the same considered feature groups (Figure 4A). Conversely, at 8.5 weeks the best differentiation in whether dopamine was released from the DLS or DMS is achieved with the RF algorithm. The aggregate accuracy rates with RF exceed 0.6, with an accuracy rate of 0.69 noted when using all 8 statistical features. Conversely, at 8.5 weeks, the SVM performance with the combined features is only marginally above chance with an aggregate accuracy rate of 0.568. For this latter scenario, the machine trained on nIRCat recordings classifies the majority of dopamine release events as having arisen from the DMS (Figure 4B). Thus, a rather high accuracy rate of 0.798 is noted in differentiating DMS recordings, but poor aggregate accuracy is apparent via the classification of the majority of DLS dopamine releases having stemmed from the DMS. Explicitly, out of the 1000 iterations of Figure 1, 798 iterations were accurately classified as having been stemmed via DMS stimulation, whereas only 318 iterations were accurately classified as having resulted from DLS stimulation. At 12 weeks the SVM again yields superior predictive capability over RF with accuracy rates of 0.64 with the 10 (i.e. combined) features, and 0.708 with 8 statistical features (Figure 4C). At each of the three animal ages, the use of the paroxysmal features does not provide above chance predictive accuracy of enabling differentiation of the signal as having originated from DLS or DMS. In general, higher accuracy rates are noted in Figure 4 when considering the statistical or combined features rather than paroxysmal features (not pictured). The better accuracies obtained with statistical or combined features highlights the importance of considering statistical rather than transient or paroxysmal aspects of the dopamine recordings for the purpose of identifying the striatal region from which those dopamine modulatory features originate: whether the recorded image is from the DLS or DMS at each of the three timepoints. In summary, at the three considered animal ages, when providing a 0.3 mA stimulation pulse to elicit dopamine release, nIRCat dopamine images and their features do provide a biomarker for differentiating whether the dopamine was released from the DLS or DMS region. Interestingly, the predictive capability in differentiating the brain region with 0.3 mA electrical stimulation increased as a function of animal age, similarly to our age-dependent ability to use our machine learning algorithms to distinguish between stimulation strengths. Specifically, the best aggregate accuracy rate at 4 weeks was 0.629 while that at 8.5 and 12 weeks were 0.69 and 0.708, respectively. These results support our prior findings and hypothesis that early in animal development there exists high variability in dopamine release and reuptake activity in a manner that equilibrates in adulthood.

**Figure 4.**
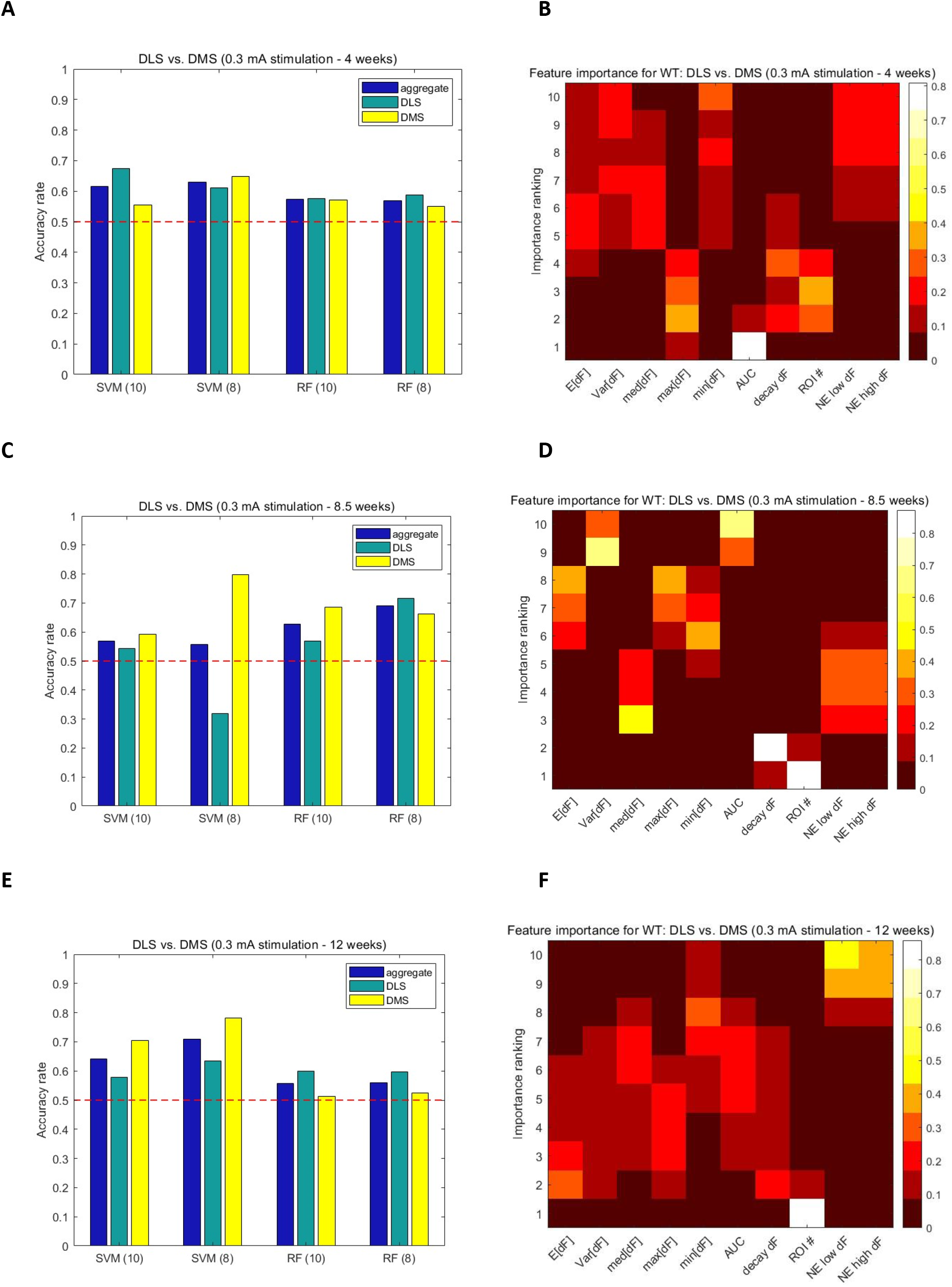
The predictive accuracy and feature importance rankings attained with the RF machine learning pipeline. Barplots quantifying RF accuracy at predicting whether the recording was made from the DLS or DMS at a stimulation strength of 0.3 mA. The quantities in the parentheses indicate the number of features that were used to train the machine and form the test data in the machine learning analysis. The subfigures (A), (C), and (E) correspond to results for the data collected from animals of 4, 8.5, and 12 weeks of age, respectively. The corresponding heatmaps (B), (D), and (F) illustrate a ranking of each feature’s importance in contributing to the RF classification decision when 10 features (Table 2) were used.

### Feature importance for differentiating DLS from DMS with dopamine modulatory signatures

We have confirmed that stimulated dopamine release imaging in acute slices provides machine learning algorithms, SVM and RF, the ability to distinguish between stimulation strength (0.1 mA vs 0.3 mA) and brain region (DLS, DMS). Interestingly, we also find that our predictive capabilities are directly proportional to animal age, supporting the hypothesis that dopamine signaling dynamics are more variable early in development and stabilize in adulthood. We next determined which features of our dopamine recordings are most important for the predictions that were made by the machine learning analyses. With RF, the node purity measure was used to determine the feature importance. The heatmaps in Figure 4(B), (D), and (F) provide an account of the frequency at which each of the 10 considered dopamine recording features were ranked in their importance for arriving at a prediction. For striatal slices obtained from mice that were 4 weeks old, the AUC was the most important feature for determining whether a recording was made from the DLS or DMS with an accuracy rate of 0.809. At 4 weeks of mouse age, the max[dF], ROI number, and decay dF were the prominent second-most important features with frequencies of 0.365, 0.284, and 0.167, respectively. Conversely, min[dF], Var[dF], and the two paroxysmal features were the least important features for distinguishing DLS from DMS at 4 weeks. At 8.5 weeks, the ROI number (frequency of 0.873) and decay dF (frequency of 0.857) were the most and second-most important features for distinguishing striatal brain region, respectively. In contrast to the 4 week time-point, the AUC was among the least important features to distinguish brain region at 8.5 weeks, while Var[dF] was again deemed a feature unimportant to distinguish brain region, consistent with our 4 week analysis. At 12 weeks, ROI number is the most important feature to distinguish DLS From DMS with the highest frequency (0.859) while the two paroxysmal features and min[dF] are the least important features in the RF determination of the brain region that had stimulation-induced dopamine release measured by nIRCat. In sum, max[dF] and ROI number - representative of maximum amount of dopamine released and number of dopamine release sites, respectively - were consistently the most important features to distinguish DLS from DMS striatal brain regions. Conversely, features associated with the variability of the baseline nIRCat nanosensor fluorescence such as Var[dF] and min [dF] were consistently unimportant in distinguishing brain regions, as expected.

A strength of inference and prediction via machine learning lies in evaluating a multidimensional dataset where a single predictive feature might not exist. Machine learning approaches can also facilitate a blind analysis of a dataset to avoid evaluator bias. Our machine learning algorithm highlighted max[dF] and ROI number as features that are consistently important across timepoints for distinguishing DLS from DMS brain regions. To verify that these features enable discrimination of nIRCat images taken from DLS versus DLS, we proceeded with a methodological analysis of our datasets, considering max[dF] and ROI number. We plotted the distribution of these features to visualize the difference in recorded values between DMS and DLS at each time point (Figure 5, Figure S1). In Figure S2, we analyzed max[dF] and ROI number values for 0.3 mA stimulation strengths across all three timepoints, and as a control, analyzed Var[dF] for 0.3 mA stimulation strengths across all three timepoints. Specifically, a two-sample Kolmogorov–Smirnov (KS) test was performed to determine whether the calculated features vary statistically significantly between the nIRCat recordings collected from the DLS and DMS. Interestingly, the only feature that met the significance criterion was the AUC when the recordings were made at 4 weeks. This corroborates the utility of the ML analysis in distinguishing differences that would otherwise not be identified as significant with conventional, univariate statistical techniques.

**Figure 5.**
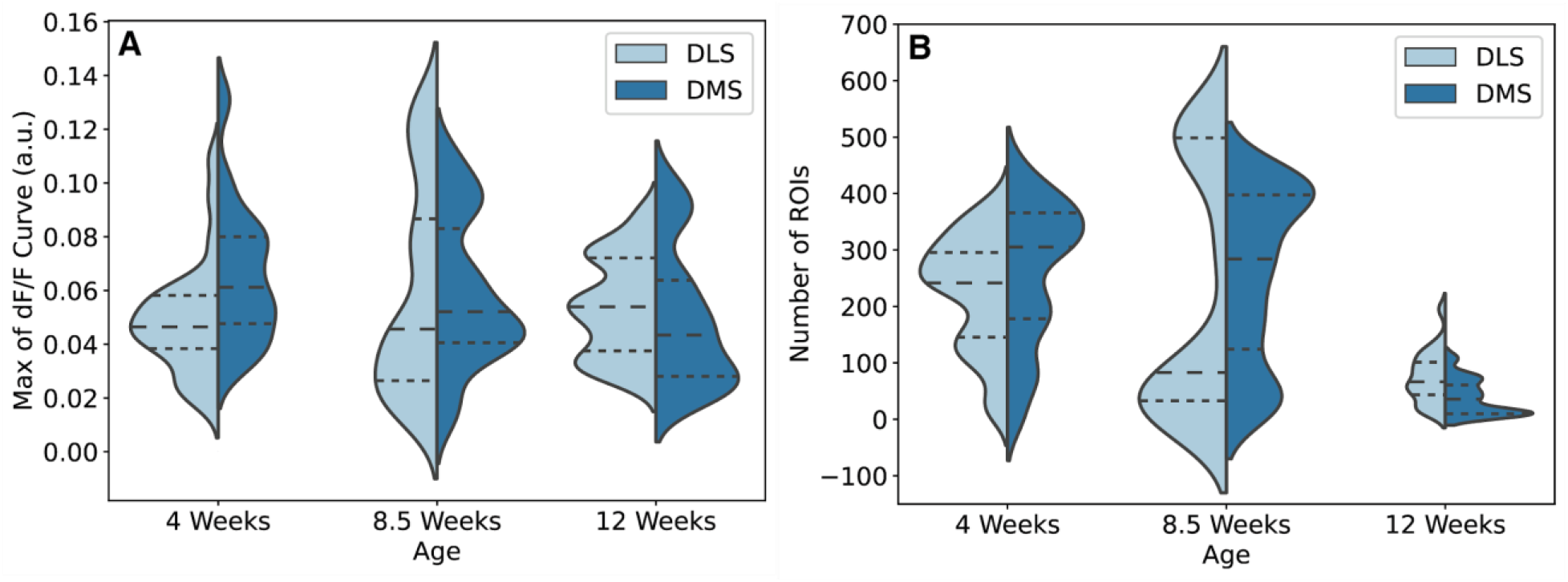
Univariate distribution of several features deemed important for distinguishing DLS from DMS brain regions. Violin plots of max[dF] (A) and ROI number (B) across age groups and DLS (light blue) versus DMS (dark blue) brain regions. Dashed lines indicate the 25th, 50th and 75th percentiles of the distribution.

### Conclusions

The past few years have seen the development of various tools to image neuromodulators, and specifically dopamine modulatory dynamics, at the spatiotemporal scales of relevance for endogenous neurochemical signaling. As these tools emerge, the rich datasets they generate in the form of dopamine signaling dynamics videos provide many features, some of which may be more helpful than others in distinguishing biological phenomena. It is therefore of interest to develop and assess tools that quantify the release, diffusion, and reuptake of neuromodulators such as dopamine, where the spatial and temporal dynamics observed is dependent on dopamine receptor activation, dopamine transporter activity, neuronal activity, and neuronal tissue microstructure. It is also important to develop computational analyses, particularly those that remove datasets from manual analysis to reduce bias.

In this manuscript, using two standard machine learning techniques with features computed via dopamine nIRCat recordings, we are able to distinguish between striatal subregions and stimulation strengths in adult mice using a support vector machine and a random forest classifier. The latter addresses the bias-variance trade-off by reducing variance and avoiding overfitting, while the former provides robust performance.

The results indicate that an accurate discernment of the stimulation strength in the DLS and DMS is not possible from very young mice (i.e. prepubescent or young adult), however, as mice age the distinguishability increases in both brain regions. The prediction accuracy of our machine learning algorithms consistently increase as a function of animal age. Increased variability in the kinetics and features of dopamine release in young animals is noted, and likely to contribute to this reduced accuracy in distinguishability. Our findings suggest that by the time animals reach adulthood (i.e. 12 weeks), it is possible that dopamine release and reuptake features have stabilized thus enabling more accurate distinguishability among the stimulation strengths.

Overall, higher predictive capabilities were noted with the statistical or combined features suggesting that entire-recording statistics rather than transient or paroxysmal aspects of dopamine modulatory kinetics are biomarkers of the striatal region from which those dopamine signals originated. It is interesting that, similar to the age-dependent increase in the capability to distinguish among stimulation strengths, the differentiability of a brain region increased with the age of the animals. The results indicate that dopamine release and reuptake dynamics are rather variable early in animal development but equilibrate in adulthood.

Future advancements to this work are possible on the computational front. It would be advantageous to use more sophisticated kernel functions that are matched to the properties of the waveforms. This would involve deriving and incorporating a priori information from dopamine signals and the experimental conditions. The incorporation of additional features also constitutes an extension, such as features that account for the timing properties of the waveforms such as *τ*_off_.

More complex classification tasks or combining different neuronal activity data types could enhance accuracy at making correct predictions with neurochemistry datasets. For instance, adding more features could enhance the translation of all traces further enhancing our classification ability. Similarly, merging datasets from both neurochemical efflux and neuronal activity could further enhance the predictive capabilities of our algorithm. Nonetheless, our current work demonstrates that machine learning approaches are able to differentiate DLS from DMS striatal sub-regions of the mouse brain, in a manner unachievable with statistical analysis of the data alone. These findings suggest machine learning approaches to studying neurochemical dynamics could help differentiate between cohorts when classical statistical analyses find no differences with single-feature comparisons. For instance, recent work has shown that dopamine dynamics in late stage Huntington’s Disease show a blunted ability to release dopamine at the single synapse level, and suggest dopamine dysregulation is driven by D2-autoreceptor regulation of dopamine release through Kv1.2 channels in late stage Huntington’s Disease (17). However, these findings were only shown to be statistically significant for late-stage Huntington’s Disease, despite trends at earlier disease timepoints that suggested dopamine signaling deficits present earlier in disease. Machine learning tools could help confirm the absence or presence of differences in such disease cohorts, potentially enabling pinpointing earlier onsets of variabilities in neurochemical signaling and what features of signaling drive those differences.

Herein, we have developed our machine learning algorithms and demonstrated their capabilities in differentiating datasets in a manner unachievable with statistical analysis of single features. However, the methods evoked in this work are not specific to dopamine signaling nor nIRCat-acquired data, rather they could be extrapolated to other time series data such as other neuromodulator probes emerging in the neuroimaging toolkit (2), (3), (4), or imaging datasets based on neuron activity such as GCaMP-based probes. Expansion of this work could follow the same workflow and may identify previously unnoticed trends. Moreover, future work may be able to identify and introduce new features that may raise the classification ability of a classifier. An exciting future avenue is to consider whether above-mentioned different neurochemical or neuron activity datasets can collectively improve the precision of machine learning algorithms when trained on these different neuronal signals. It would be interesting to investigate whether the same features we identified in dopamine modulatory traces (i.e. peak dF/F and ROI number) are the most predictive for other neurotransmitters that exhibit different signaling dynamics in different brain regions. It is possible that separate machines must be trained to make predictions when considering different neurotransmitters in parallel, or that independent datasets would collectively enhance the performance of a machine, despite data inputs from different biological pathways. Perhaps most interesting to consider is that many of these datasets already exist in the literature and could readily be tested for feature mining with machine learning approaches for a more holistic and userin-dependent approach to data analysis in neuroimaging.

## Methods

### nIRCat sensor production

nIRCat nanosensor is produced by combining 100 μL of a 2 mg/mL solution of HiPCO raw single walled carbon nanotubes (SWNT) suspended in molecular biology grade water, 100 μL of a 1X phosphate buffered saline (PBS) solution, 100 μL of a 1mM (GT)_6_ ssDNA solution. The combined solution is probe tip sonicated for 10 minutes then centrifuged at 16,000g for 30 minutes. The supernatant is then filtered through a 160k Da spin filter for 5 minutes at 8,000g. The retentate is then resuspended in water and centrifuged for 5 minutes at 1,000g. The resulting solution is a fully prepared nIRCat sensor. The sensor concentration is then calculated by using absorbance measured at 632 nm. Dopamine response is confirmed by measuring the fluorescence response of 2 mg/mL solutions of the sensor before and after addition of dopamine (2), (17), (18), (19).

### Plant acute brain slice generation and sensor labeling

All procedures involving animals were approved by the University of California Berkeley Animal Care and Use Committee. Both male and female B6CBAF1/J mice (https://www.jax.org/strain/100011) were used for experiments. Mice were group-housed after weaning on P\postnatal day 21 (P21) with nesting material on a 12:12 light cycle. Acute brain slices were produced from three age groups: 4 weeks (P32 - P35), 9.5 weeks (P64 - P66), and 12 weeks (P87 - P92). To generate acute brain slices, mice were anesthetized via intraperitoneal injection of a ketamine/xylazine cocktail. Mice were perfused transcardially using chilled, ascorbic acid free cutting buffer (119 mM NaCl, 26.2 mM NaHCO3, 2.5 mM KCl, 1 mM NaH2PO4, 3.5 mM MgCl2, 10 mM glucose, and 0 mM CaCl2). The brain was extracted and mounted on a vibratome cutting stage (Leica VT1200 S) to produce 300 μm coronal slices containing the dorsal striatum. Slices were left in ACSF (119 mM NaCl, 26.2 mM NaHCO3, 2.5 mM KCl, 1 mM NaH2PO4, 1.3 mM MgCl2, 10 mM glucose, and 2 mM CaCl2) to rest at 37°C for 30 minutes followed by 30 minutes at room temperature. Slices were labeled with nanosensor by transferring the slices to a small volume incubation chamber containing oxygen-saturated ACSF and adding nIRCat nanosensor for a final concentration of 2mg/L. Slices were subsequently rinsed in ACSF by transferring the slices through three wells of a 24-well plate and left to rest in ACSF for 15 minutes. Labeled slices were then transferred to the microscope recording chamber and allowed to equilibrate for 10 minutes before imaging. All imaging experiments were performed at 32°C.

### Stimulation and image collection

Following incubation with nIRCats, slices are placed underneath a near-Infrared optical microscope (2). Specific brain regions were identified using a 4x objective (Olympus XLFluor 4×/340). A bipolar stimulation electrode was placed 200um away from the region to be imaged. Using a 60x objective, 600 images were collected at a frame rate of 8.33 Hz with a 100ms stimulation occurring at the 200th frame. This process was repeated until the desired amount of replicates was achieved. During this process, slices were kept viable by flowing an oxygenated ACSF solution held at 34°C over the slices at a rate of 2mL/min.

### Image processing

Image stacks were imported into MATLAB and were segregated into a grid consisting of 25 by 25 pixels (or roughly 7 by 7 microns). Each box within the grid was considered a region of interest (ROI). A moving average was used to calculate the baseline of each ROI and subtracted from the original trace. The resulting trace left for the ROI was a time series of change in fluorescence. Between frames 200 and 300, the algorithm looked for an event that was 3 standard deviations above the average noise previously calculated for this ROI. If an event was found, this ROI was marked significant. If no event was detected, this ROI was deemed inactive and no more calculations are completed on this ROI. For each significant ROI, two exponentials were fit to the trace to determine the time constant of turn on and turn off. The algorithm also finds values such as maximum change in fluorescence and area under the curve (2).

### Machine learning classification methods

The SVM and RF algorithms were trained on the features in Table 2 to differentiate among the stimulation strength or the location of dopamine release as described above. The SVM used a linear kernel with a binary classifier and was implemented in R via the packages e1071 and kernlab (13). A linear kernel was used for simplicity and also the lack of a priori knowledge of the relationships between the features and the dynamics and source of dopamine release. The RF was implemented via the randomForest function and package in R (14). The number of variables used for each split in the tree was specified as 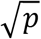 with p=10 or p=8 for the scenarios of using the combined or statistical features, respectively. In all cases, the number of trees used was 1000. We did not consider RF with the paroxysmal features as there were too few variables (i.e p=2) to consider this as worthwhile analysis. The evaluation of the predictive capability via the classifier accuracy rate entailed a leave-one-out analysis with Monte-Carlo (MC) sampling of the brain slices. The MC analysis consists of the data being repeatedly divided into a training and test set with the test data consisting of a single observation while the remaining data was evenly partitioned into two groups and used to train the machine. The classification decisions that consisted of distinguishing between possible stimulation strength or brain regions were binary. For each scenario the accuracy rates for the two possibilities were presented in the figures. The third consideration deemed the “aggregate” consisted of averaging the possibilities together to provide a holistic account. Our consideration of feature importance consisted of the node purity metric in the RF technique. The metric was used to evaluate which of the stimulated dopamine features were the most important in the classifiers’ decisions of the brain region or stimulation strength.

## Data and Code availability

Data and code are available upon request.

## Acknowledgements

We acknowledge support of a Burroughs Wellcome Fund Career Award at the Scientific Interface (CASI) (MPL), a Dreyfus foundation award (MPL), the Philomathia foundation (MPL), an NIH MIRA award R35GM128922 (MPL), an NIH R21 NIDA award 1R03DA052810 (MPL), an NSF CAREER award 2046159 (MPL), an NSF CBET award 1733575 (to MPL), a CZI imaging award (MPL), a Sloan Foundation Award (MPL), a USDA BBT EAGER award (MPL), a Moore Foundation Award (MPL), and a DOE office of Science grant DE-SC0020366 (MPL). MPL is a Chan Zuckerberg Biohub investigator, a Hellen Wills Neuroscience Institute Investigator, and an IGI Investigator.

