## Supplemental Information for "Identifying Neural Signatures of Dopamine Signaling with Machine Learning"

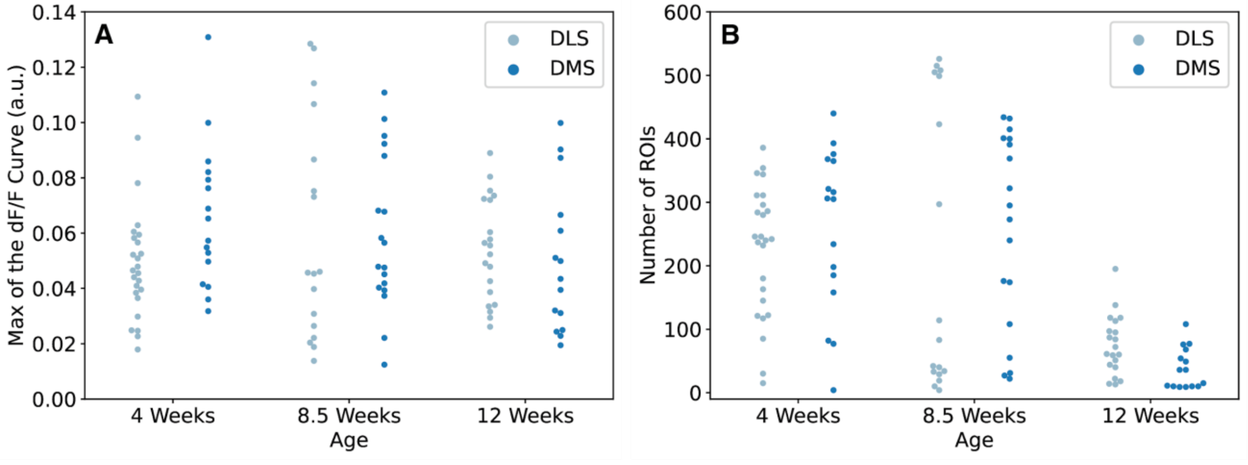


**Figure S1. A visualization of the difference in recorded values between DMS and DLS at different time points in mouse development.** Swarm plots of the features (A) max[dF], and (B) ROI number across age groups and DLS (light blue) versus DMS (dark blue) brain regions. Each point indicates a measurement from a single recording.

**A**

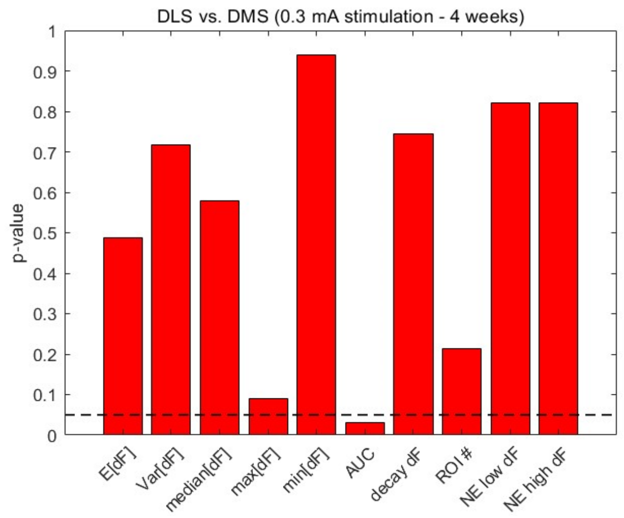

**B**

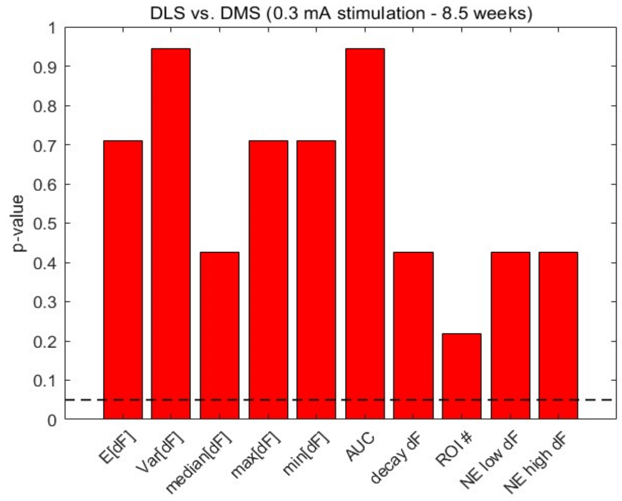


**C**

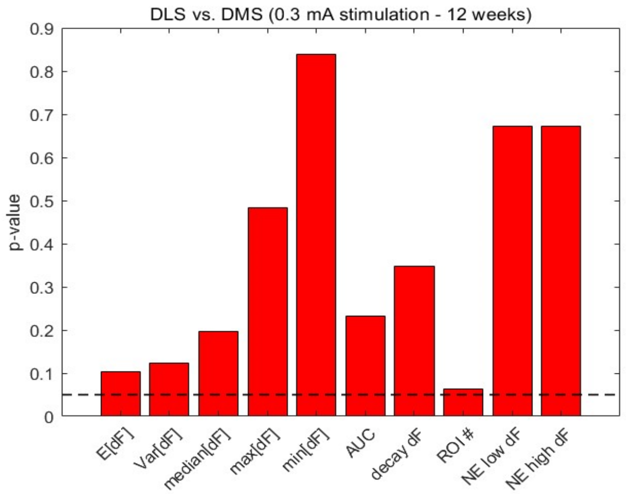


**Figure S2. Univariate statistical analysis of calculated features from the DLS and DMS for 0.3 mA stimulation strengths across all three age groups.** The p-values resulting from two-sample Kolmogorov-Smirnov (KS) tests applied to the 8 statistical features and 2 paroxysmal features obtained from recordings at (A) 4, (B) 8.5, and (C) 12 weeks in differentiating whether the DMS or DLS was stimulated at 0.3 mA. The p-values below the dashed, black line reflect the rejection of the null-hypothesis at $p < 0.05$ that the values attained for the feature arose from the same distribution irrespective of the region stimulated. The only feature that met this criterion was the AUC when the recordings were made at 4 weeks.
